# Exogenous Nitro-Oleic Acid inhibits primary root growth by reducing the mitosis in the meristem in *Arabidopsis thaliana*

**DOI:** 10.1101/2020.06.17.155416

**Authors:** Luciano M. Di Fino, Ignacio Cerrudo, Sonia R. Salvatore, Francisco J. Schopfer, Carlos García-Mata, Ana M. Laxalt

## Abstract

Nitric oxide (NO) is a second messenger that regulates a broad range of physiological processes in plants. NO-derived molecules called reactive nitrogen species (RNS) can react with unsaturated fatty acids generating nitrated fatty acids (NO_2_-FA). NO_2_-FA work as signaling molecules in mammals where production and targets have been described under different stress conditions. Recently, NO_2_-FAs were detected in plants, however their role(s) on plant physiological processes is still poorly known. Here we show that exogenous application of nitro-oleic acid (NO_2_-OA) inhibits Arabidopsis primary root growth; this inhibition is not likely due to nitric oxide (NO) production or impaired auxin or cytokinin root responses. Deep analyses showed that roots incubated with NO_2_-OA had a lower cell number in the division area. Although this NO_2_-FA did not affect the signaling mechanisms maintaining the stem cell niche, plants incubated with NO_2_-OA showed a reduction of cell division in the meristematic area. Therefore, this work shows that NO_2_-OA inhibits mitotic processes subsequently reducing primary root growth.

## INTRODUCTION

The formation of nitrolipids was initially proposed in animals from the observation that nitric oxide (NO) inhibited lipid oxidation propagation reactions (Rubbo et al., 1994). The nitration of fatty acids is induced by species derived from NO (Freeman et al., 2008). In animals, nitrated lipids are signaling molecules, acting as intermediaries in potent cascades of signal transduction, which translate into changes in protein functionality due to post-translational modifications (Batthyany et al., 2006; Rubbo and Radi 2008; Trostchansky and Rubbo 2008; Schopfer et al., 2011). So far, there are few reports about the detection of NO_2_-FA in plants. Specifically, adducts between NO_2_-OA and Cys (NO_2_-OA-Cys) together with nitro-conjugated linoleic acid (NO_2_-cLA) were detected in olive fruits and Extra Virgin Olive Oil (EVOO) although nor free NO_2_-OA was detected in olives (Fazzari et al., 2014). NO_2_-OA was recently detected in the free fatty acid fraction of seeds and seedlings of *Brassica napus* (Vollar et al., 2020). Also, nitro-linolenic acid (NO_2_-Ln) was observed in cell suspension cultures, seeds, seedlings and leaves of the model plant Arabidopsis (Mata-Perez et al., 2016a) and in important crops such as rice and pea (Mata-Perez et al., 2016b). Exogenous application of NO_2_-Ln induced both and antioxidant response and the chaperone network in Arabidopsis (Mata Perez et al., 2016a) and NO production in Arabidopsis primary root tip (Mata Perez et al., 2016c). Exogenous application of nitro-oleic acid (NO_2_-OA) induced reactive oxygen species (ROS) production via activation of NADPH oxidases and not NO production in tomato cell suspensions (Arruebarrena Di Palama et al., 2020). Arabidopsis NADPH oxidase mutants showed that NADPH isoform D (RBOHD) was required for NO_2_-OA-induced ROS production in leaves (Arruebarrena Di Palma et al., 2020).

NO is a well-established second messenger in plants and is involved in the development of the root system. NO regulates processes such as the formation of root hairs in Arabidopsis and lettuce (*Lactuca sativa L*) (Lombardo et al., 2006), as well as the formation of adventitious roots via the activation of MAPK in cucumber (*Cucumis sativus*) (Pagnussat et al., 2004). The root is divided in the mersitematic/division zone, the elongation zone and the differentiation zone. The meristematic cells, a cell type with a high rate of cell division, coordinates the growth and development of the root. In Arabidopsis, NO targets the cells in the elongation zone, inhibiting the cellular elongation mediated by gibberellins (Fernandez-Marcos et al., 2012). In addition, NO decreases the levels of the auxin transporters (PINs) in the membrane, affecting the transport and distribution of auxins and altering downstream signaling (Fernandez-Marcos et al., 2011). High levels of NO inhibit the development of the primary root, reducing the meristematic zone, directly affecting mitotically active cells (Fernandez-Marcos et al., 2011).

Most of the plant hormones (abscisic acid, auxins, cytokinins, ethylene, gibberellins and brassinosteroids) regulate cell division and elongation processes (Benkova and Hejatko 2009, Wolters and Jürgens 2009). The root stem cell niche at the root apical meristem is composed of stem cells with high rate of division given rise to specific root cells lineages, which are surrounding a group of cells with a low division rate called the quiescent center (QC) (Scheres 2007). QC plays an important role in maintaining undifferentiated stem cells (van den Berg et al., 1997). The differentiation of daughters stem cells give rise to several cell types integrating the root architecture, being growth defined by the balance between cell division and elongation (Dolan et al., 1993; Scheres et al., 1994).

In this work we show that NO_2_-OA inhibits primary root growth. This inhibition is not due to NO signaling, altered auxine/cytokinin responses or altered mechanism that maintain the stem cell niche. Here we show that exogenous application of NO_2_-OA reduce the cell cycle marker CYCB1:1, resulting in a short meristem.

## RESULTS

### NO_2_-OA inhibits primary root growth in Arabidopsis

Arabidopsis seedlings present basal levels of NO_2_-Ln that increase when seedlings are subjected to wounding, cadmium or low temperature stress (Mata Perez et al., 2016a). All major nitro lipids share the same electrophilic center, similar reactivity properties and therefore a common mechanism of action (Baker *et al.*, 2004). In animals, NO_2_-OA has long been used as a surrogate to study and understand the regulation, signaling and metabolism of nitrated fatty acids given its additional stability and well developed synthetic routes (Freeman et al., 2008). We studied the effect of exogenous application of NO_2_-OA on primary root growth of Arabidopsis seedlings. **Figure 1** shows dose-dependent inhibition of primary root growth upon NO_2_-OA treatments compared to untreated or oleic acid (OA) treated seedlings.

**Figure 1:**
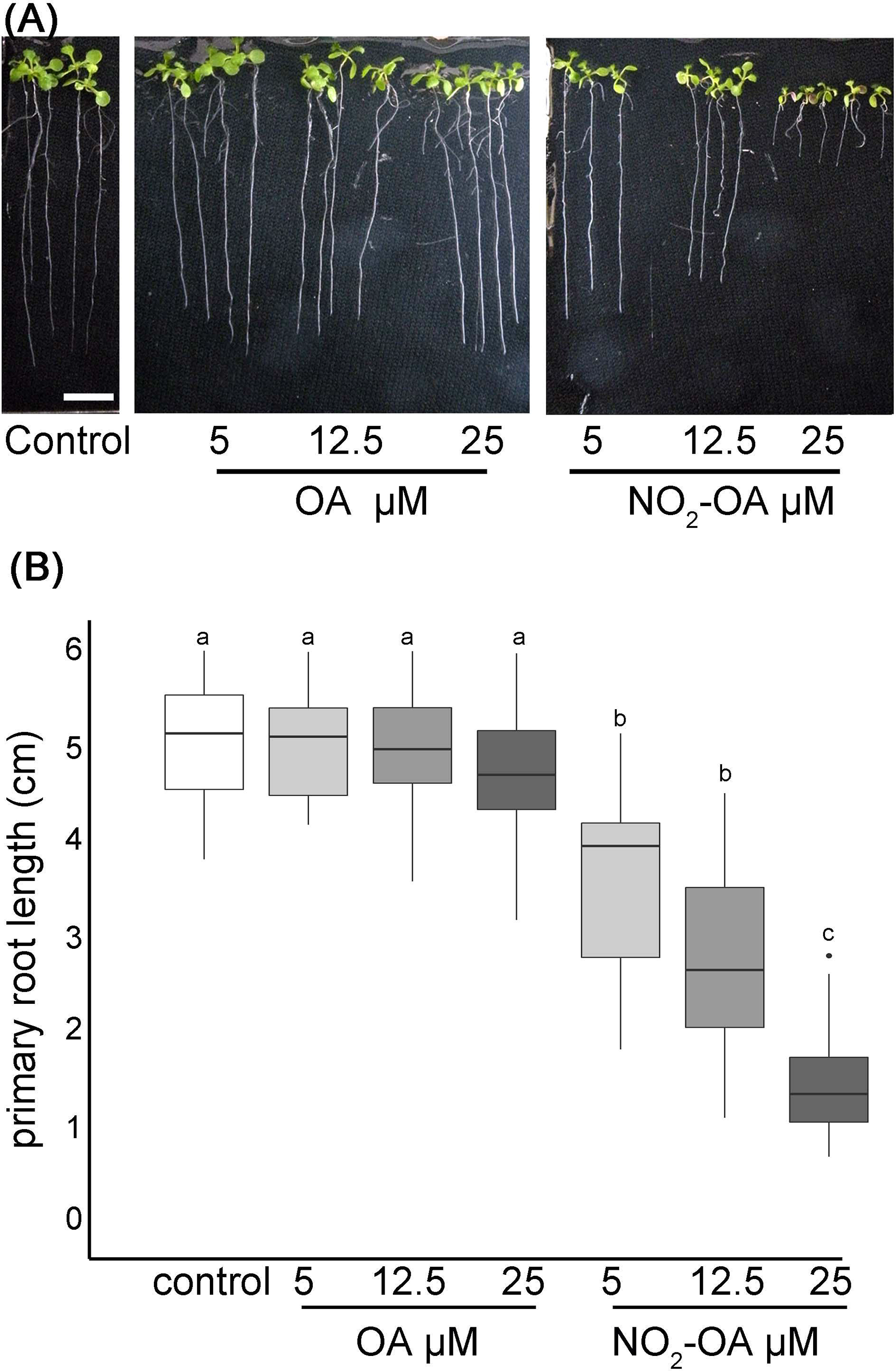
NO_2_-OA inhibits primary root growth in Arabidopsis. Seeds were germinated and vertically grown in MS agar for five days, then transferred onto plates with NO_2_-OA or OA (25, 12.5 o 5 μM) during five days. **(A)** Representative image of Arabidopsis primary root growth. Scale bar = 1 cm **(B)** Primary root length quantification. Data is depicted in box-plot graphs were the box is bound by the 25th to 75th percentile, whiskers span to minimum and maximum values, and the middle line represents the average of 5 independent experiments. Different letters indicated statistical significant differences, n=25 (ANOVA, Tukey p<0.001).

The inhibition of the primary root growth could be due to a lower number of cells at the apical meristem and/or a lower cell size or cell elongation rate on the elongation zone of the root. To determine the processes by which NO_2_-OA is modulating root growth, we measured the size and the number of root cells in the meristem. Five-day old Arabidopsis seedlings were treated for another five days with NO_2_-OA or OA, and the root cells were analyzed by DIC microscopy. The column of cortex cells was counted from the quiescent center (QC) to the last cell where its length was not greater than 50% compared to the previous one according to what was described in Perilli & Sabatini 2010 **(Figure 2 A)**. As shown in **Figure 2 B** control or OA treated roots have an average of 36±5 cortex cells in the division zone. NO_2_-OA-treated roots have a significant reduction in the number of cortex cells **(Figure 2 B)**. Roots treated with 12.5 or 25 μM of NO_2_-OA showed an average of 29±4 and 19±4 cells, respectively. The extension of the meristematic zone was measured as the distance from the QC to the last cell of the division zone **(Figure 2 C)**. In control roots, the meristematic zone measures 300±44 μm, while NO_2_-OA treated roots shows a strong reduction of the meristematic zone (240±40 μm and 180±22 μm for 12.5 or 25 μM NO_2_-OA respectively). The cell size at the elongation zone was not significantly different between treated or not treated roots, being the size of cortical elongated cells 168.8±2.9, 171.3±2.6 and 166.7±1.9 μm (mean ± SE, n=6) for control, OA and NO_2_-OA-treated roots, respectively **(Figure 2 D)**. A establish balance between cell division and cell differentiation governs meristem size and thus root growth. Since no differences were found in the size of the cells in the differentiation zone, and a reduction of number of cortical cells is observed, then the inhibition of root growth is probably due to a regulation of cell division process.

**Figure 2:**
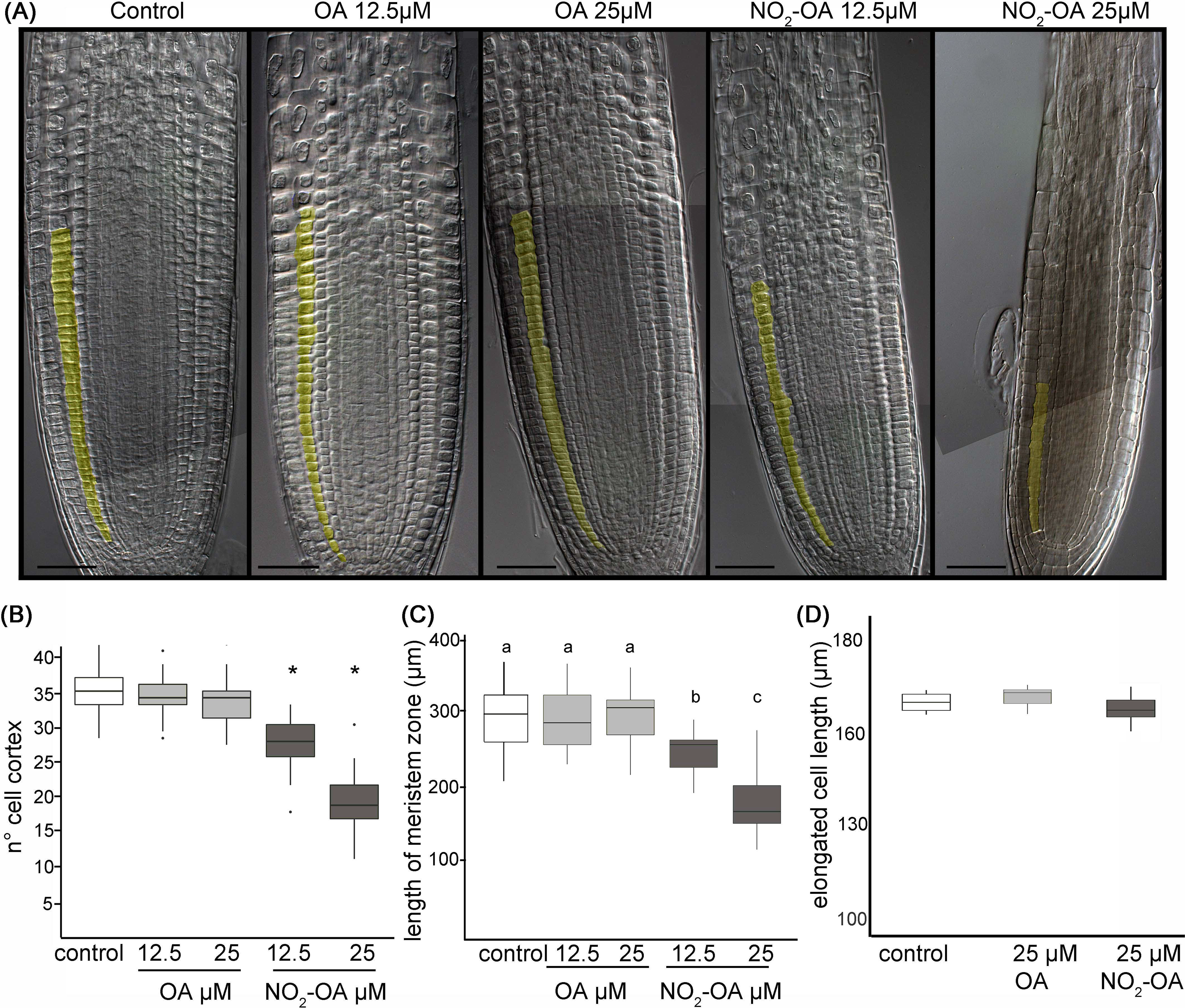
Effect of NO_2_-OA on the Arabidopsis root meristem. Five-day old seedlings were treated with NO_2_-OA or OA (25, 12.5 μM) or not treated during other five days. Roots were cleared with Hoyer’s solution and observed under DIC microscope. **(A)** Images of roots from seedlings showing cortex meristem cells artificially colored. Scale bars = 10 μm. **(B)** Meristem cell number. Asterisks indicate significant differences from control, n=30 (p<0.001, Poisson). **(C)** Size of root meristem. Different letters indicate statistically significant differences, n=30 (ANOVA, Tukey p<0.001). **(D)** Total cell size in elongation zone.

It has been previously reported that NO inhibits the growth of the primary root in Arabidopsis (Fernandez-Marcos et al., 2011). In addition, in Arabidopsis roots and cell suspensions, NO_2_-Ln treatments induced NO production (Mata-Pérez *et al*., 2016c). Therefore the evidence suggested the NO_2_-OA might be affecting root growth in a NO-dependent manner. However, the analysis of NO production, using the florescent probe DAF-FM-DA in 25 μM NO_2_-OA treated roots shows no florescence in OA or NO_2_-OA treated roots (**Figure 3**). We used 100 μM of the NO donor SNP as a positive control in order to confirm the functionality and distribution of the probe within the root. **Figure 3** show an intense and equally distribution of florescence throughout the root tip in SNP treated roots. These results show that NO_2_-OA treatment does not trigger NO production in Arabidopsis seedlings suggesting that NO_2_-OA is not inhibiting primary root growth via NO signaling.

**Figure 3:**
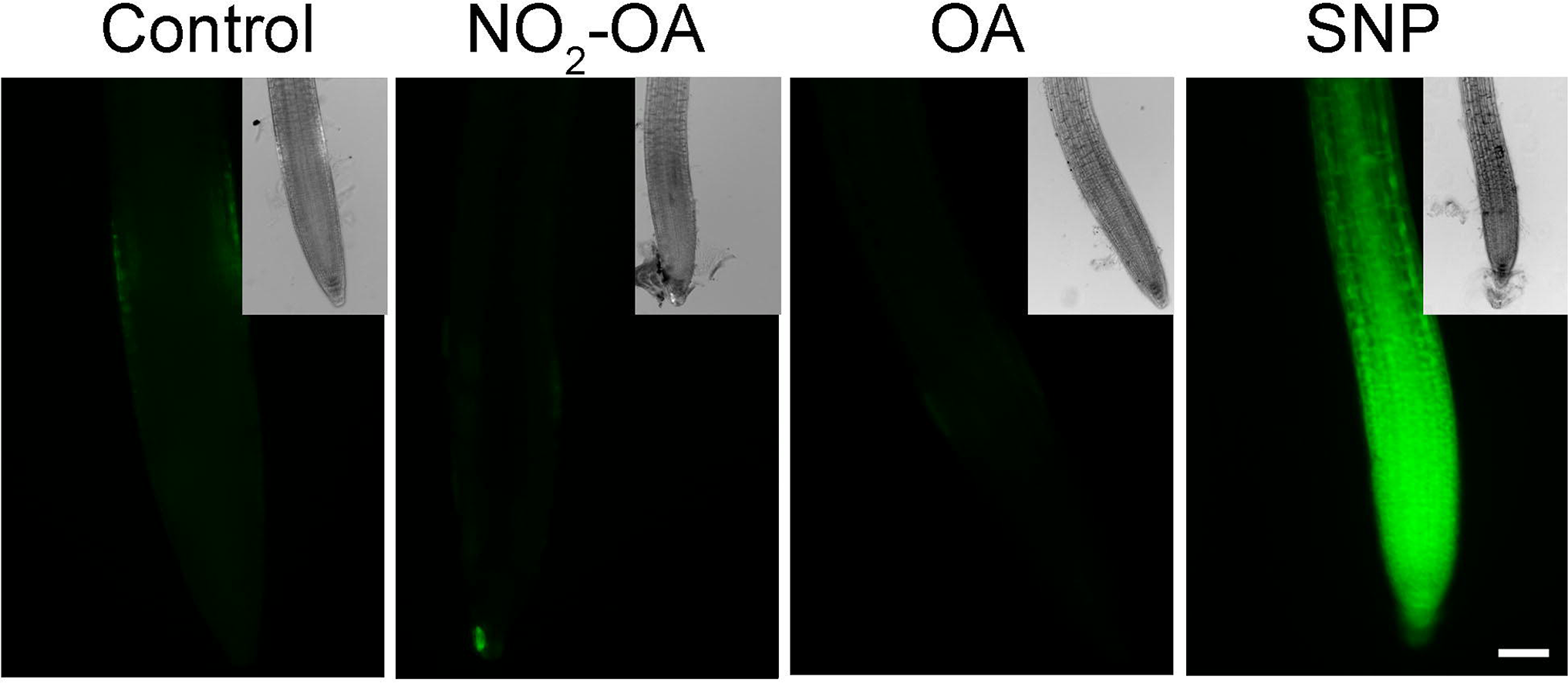
NO_2_-OA does not trigger NO accumulation on the meristematic area. Col-0 seedlings were grown for 5 days being after either not-treated or incubated with 25 μM NO_2_-OA or OA for five-extra days. Roots were incubated with DAF-FM DA 10 μM for 30 min. As positive control for DAF-FM DA probe, roots were treated with 100 μM SNP for 30 min. Scale bar= 10 μm. At least 12 roots were visualized of 3 independent experiments.

### NO_2_-OA reduces the number of mitotic cells

A reduction in the number of meristem cells could be due to a hormonal unbalance. Auxin distribution is important in regulating primary root growth. One of the described roles for auxin in the root tip is to maintain the stem cell niche and promote cell division (Grieneisen et al., 2007). In order to study auxin-response after NO_2_-OA application, we used Arabidopsis plants with auxin response reporter *DR5_pro_:GUS*. *DR5_pro_:GUS* activity is observed in the stem cell niche, columella cells and a more subdued staining in vascular tissue in all treatments. **(Figure 4 A).** Therefore, this data suggests that auxin responses are not altered because of NO_2_-OA application. In order to confirm the latter result, we used DII-Venus as an independent reporter that shows endogenous auxin abundance, reflecting the input into the auxin-signaling pathway. DII-Venus is a fusion of auxin-dependent degradation domain II of an Aux/IAA protein to Venus fluorescent protein, such that the absence of fluorescence marks auxin accumulation (Brunoud et al., 2012). In agreement with DR5pro:GUS activity, auxin levels (evidenced as the absence of fluorescence) are high in the columella, the quiescent center, and the differentiating xylem cells on control or 12,5 μM NO_2_-OA or OA treated roots **(Figure 4 B)**. In the meristem zone, from QC to TZ, auxin levels are low in control and OA treated roots, visualized as an intense fluorescence of the DII-Venus protein in the nuclei of the cortex and epidermal cells **(Figure 4 B)**. However, when we quantified the fluorescence intensity, NO_2_-OA treated roots showed less fluorescence compared to OA or non-treated roots **(Figure 4 C)**, suggesting an increase in auxin level. However, exogenous application of auxin to wild-type Arabidopsis roots caused an increase in meristem size (Dello Ioio et al., 2007), which is in contraposition to the observed effect that NO_2_-OA has on root growth. Thus, we cannot discard other explanations for a slight reduction on the fluorescence levels of DII-Venus in NO_2_-OA treated root cells, especially since DII-Venus is a semiquantitative reporter. Despite that fact, both auxin reporters, DR5pro-GUS and DII-Venus, unequivocally show that NO_2_-OA does not decrease auxin levels at the stem cells, columella or vascular tissue. Thus, auxin responses (**Figure 4 A**) and levels (**Figure 4 B and C**) are not altered during NO_2_-OA application on stem cells, columella and the differentiating vascular cells.

**Figure 4:**
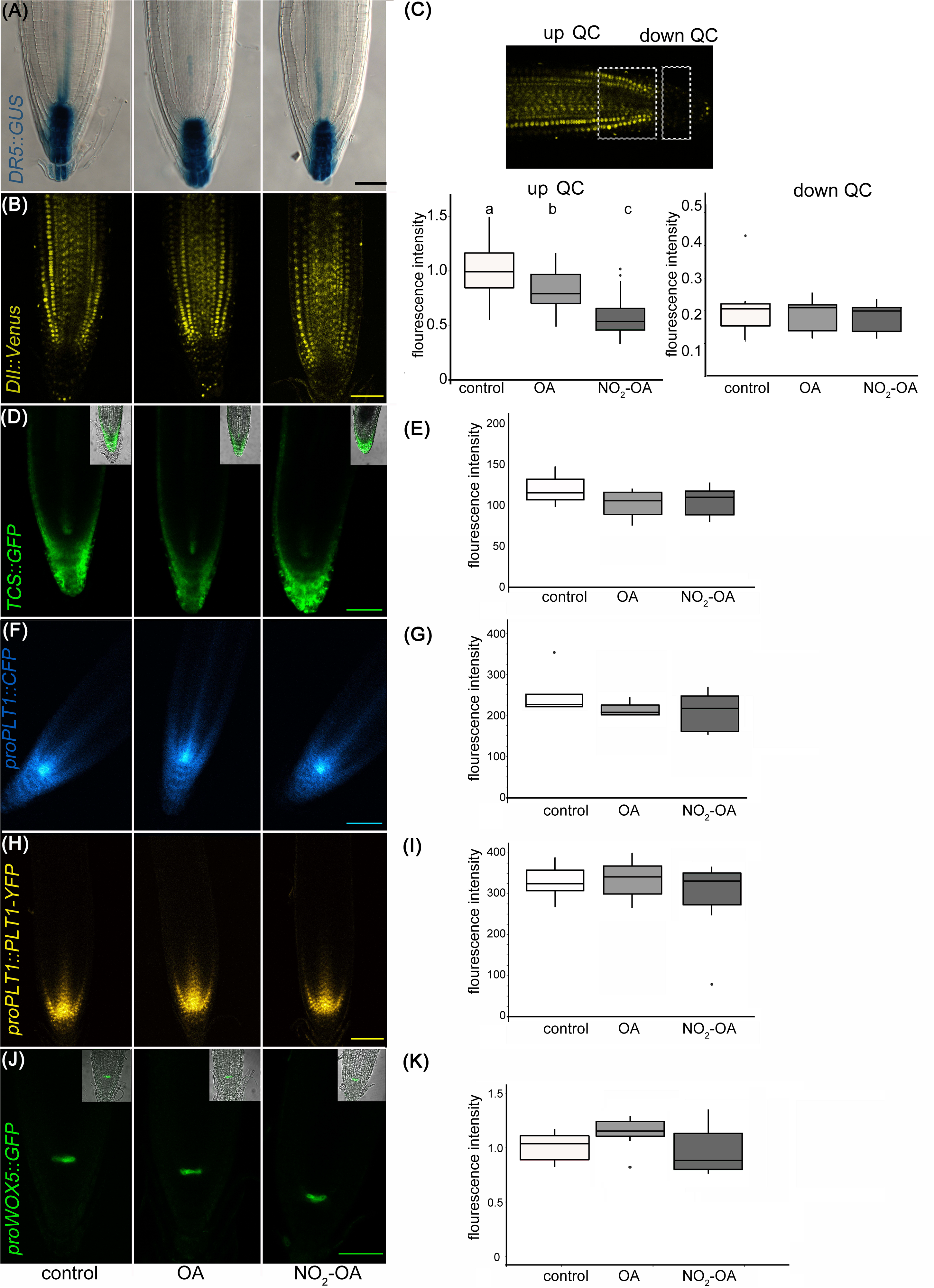
Effect of NO_2_-OA on hormonal signaling. Seedlings from different reporter lines were grown for 5 days being after either not-treated or incubated with 12.5 μM OA or NO_2_-OA for five-extra days. Representative images are shown. **(A)** Close-up view of GUS staining in *DR5_pro_:GUS* seedlings. At least 15 roots were visualized of 3 independent experiments. **(B)** Confocal images of seedlings with the *DII:Venus* reporter line. **(C)** Fluorescence intensity measured at two different root zones, up which includes de QC or down quiescent center. At least 12 roots were visualized of 3 independent experiments. Difference letters indicate statistical significant difference (one way ANOVA, post hoc Tukey p<0.05) **(D)** Confocal images of seedlings with the *TCS:GFP* reporter line. **(E)** Fluorescence intensity measured from at least 12 roots from 3 independent experiments (one way ANOVA, post hoc Tukey p<0.05). **(F)** Confocal images of seedlings with the *proPLT1::CFP* reporter line. **(G)** Fluorescence intensity measured from at least 12 roots from 3 independent experiments (one way ANOVA, post hoc Tukey p<0.05). **(H)** Confocal images of seedlings with the *proPLT1::PLT1-YFP* reporter line seedlings. **(I)** Fluorescence intensity measured from at least 12 roots from 3 independent experiments (one way ANOVA, post hoc Tukey p<0.05). **(J)** Confocal images of seedlings of *proWOX5:GFP* reporter line. **(K)** Fluorescence intensity measured from at least 10 roots from 2 independent experiments (one way ANOVA, post hoc Tukey p<0.05). Scale bars = 10 μm. The fluorescence signal intensities were quantified by using Fiji software in all the experiments. For all markers we used the same ROI (size and shape) to analyze all images of the respective experiment.

The role of auxins in root growth is associated with that of their hormone antagonists, cytokinins. It has been described that the size of root apical meristem increases in mutants from the cytokinin synthetic pathway and decreases by the exogenous application of cytokinins (Dello Ioio et al., 2007; Miyawaki et al., 2004). We studied the cytokinin response with the cytokinin reporter *TCS:GFP* using confocal microscopy (Zürcher *et al.,* 2013; Zürcher *et al.,* 2016) Under control conditions, *TCS:GFP* is observed at the root tip, particularly in the calyptra **(Figure 4 D)**. The roots treated with NO_2_-OA or OA did not show differences relative to the control **(Figure 4 E)**. Altogether the results show that the reduction of meristematic cells in NO_2_-OA treated roots is independent of auxin or cytokinin responses.

Meristematic cells come from stem cells. Stem cells are in a microenvironment, where hormone concentrations and transcription factors play a fundamental role for its maintenance. The root stem cell niches, is formed by the QC and the adjacent stem cell initials (Petricka et al., 2012), which are specified by two parallel pathways: the PLETHORA (PLT) and SHORTROOT (SHR)/SCARECROW (SCR) pathways (Petricka et al., 2012; Heyman et al., 2014). SCR maintains QC and stem cell identity (Sabatini et al., 2003), in part by inducing the expression of WUSCHEL-RELATED HOMEOBOX5 (WOX5), a QC specific gene (Sarkar et al., 2007). Two types of transcription factors have been well studied regarding the preservation of undifferentiated stem cells, PLETHORA 1 (*PLT1*, at3g20840) and PLETHORA 2 (*PLT2*, at1g51190) (Aida et al., 2004; Galinha et al., 2007; Mähönen et al., 2014). High levels of these PLT genes localize in the area of cell division, whilst low PLT levels lead root cells to expand and differentiate The presence and distribution of PLT factors in the root tip regulate the size of the meristem (Aida et al., 2004; Kornet et al., 2009; Mähönen et al., 2014). The expression of PLT1 and PLT2 has been reported to colocalize with the location of auxin in the root tip (Xu et al., 2006; Aida et al., 2004). In order to determine if NO_2_-OA affects stem cells and QC, independent transgenic lines with the transcriptional fusion reporters *proPLT1-CFP, proPLT2-CFP* or *proWOX5-GFP* and the translational fusion reporter *proPLT1:PLT-YFP* were treated with NO_2_-OA or OA. As seen in **Figure 4 F** (*PLT1* promotor activity) and **H** (PLT1 protein levels), untreated seedlings shows the highest level of *PLT1* expression in the QC and stem cells, gradually decreasing the intensity in the meristematic area. No differences were observed in this expression pattern for NO_2_-OA-treated roots **(Figure 4 G and I)**. In the case of the transcription factor *PLT2* the expression is restricted to the QC, showing no changes in any of the treatments **(supplemental figure 1)**. We analyzed the expression of QC marker *proWOX5-GFP* (Sarkar et al., 2007) and we observed the same expression pattern in NO_2_-OA compared to OA or non-treated roots **(Figure 4 J and K)**. These results show that the expression of PLT and WOX5 are not affected in roots treated with NO_2_-OA, indicating that NO_2_-FA does not affect the signaling mechanisms related to stem cell niche maintenance.

A reduction in the number of cells in the meristematic zone indicates that the process of cell division is likely to be affected, and consequently the number of cells in mitosis is reduced. To confirm this, we use transgenic plants with the transcriptional fusion reporter *proCYCB1.1-GUS* construct. *Cyclin B1.1* (CYCB1.1) is an enzyme that belongs to the cyclin family and it is expressed in the G2/M (Colón-Carmona et al., 1999). **Figure 5** shows a significant reduction on the number of cells in mitosis in NO_2_-OA treated roots with respect to control or OA treated roots, indicating the NO_2_-OA is affecting cell division pace at the meristematic zone.

**Figure 5:**
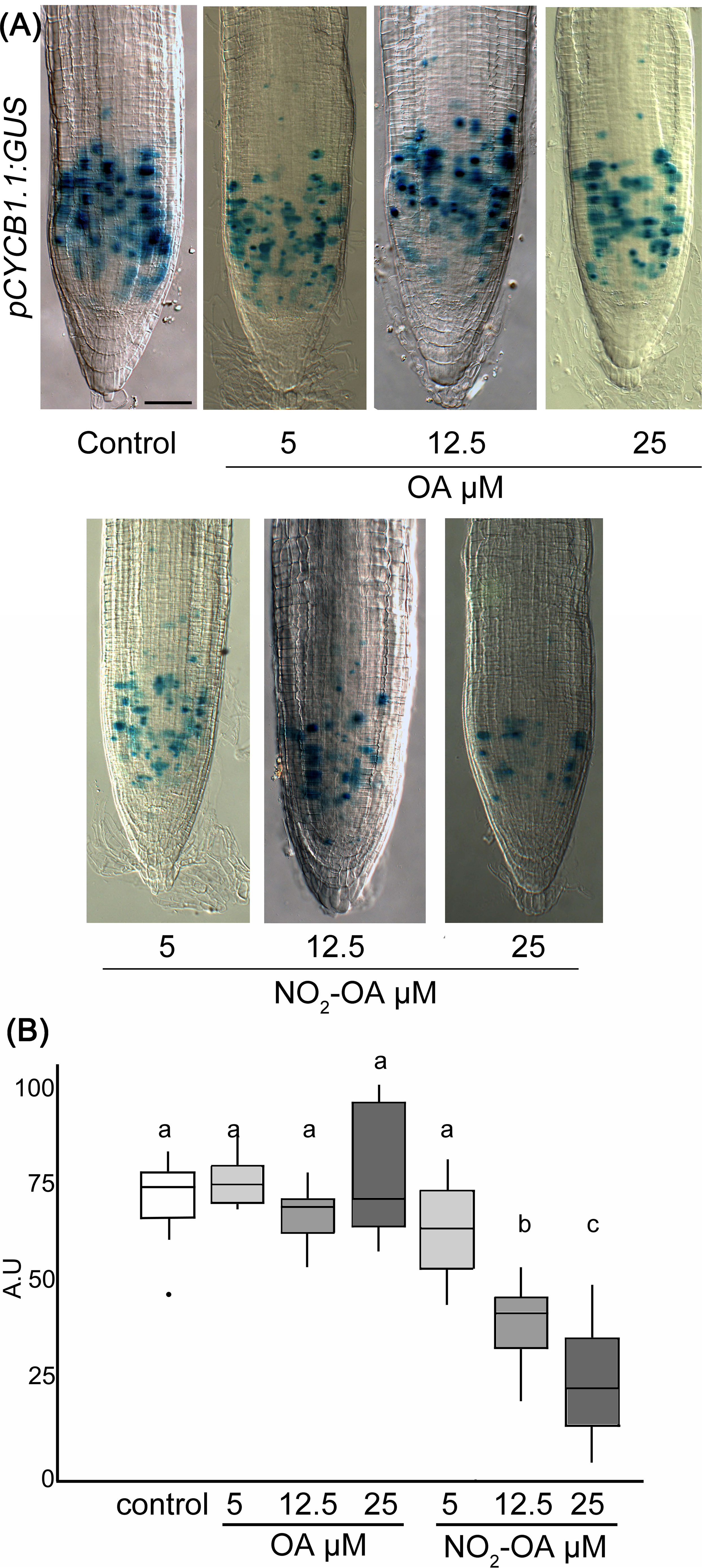
Effect of NO_2_-OA on cell-division activity. Seedlings from the pCYCB1.1:GUS reporter line were grown for 5 days being after either not-treated or incubated with 25, 12.5 and 5 μM OA or NO_2_-OA for five-extra days. **(A)** Representative close-up views of GUS staining are shown. Scale bars = 10 μm. **(B)** GUS staining quantification, difference letters indicated statistical significant difference, 3 independent experiments, n=12-15 (ANOVA, Tukey p<0.05).

## DISCUSSION

Nitrated fatty acids are well-described signaling molecules in animals (Freeman et al., 2008; Batthyany et al., 2006; Rubbo and Radi 2008; Trostchansky and Rubbo 2008; Schopfer et al., 2011). Detection of NO_2_-FA in plants was reported in Arabidopsis, pea, rice, and olives (Fazzari et al., 2014; Mata-Perez et al., 2016a,b). Our study focused on evaluating the role of a prototypical NO_2_-FA on Arabidopsis thaliana root development. For that, we decided to emulate the approach used in mammals by using the exogenous application of NO_2_-OA, allowing us to reduce the complexity of the study and focus on its functional aspects. As mentioned earlier, in animals, NO_2_-OA has long been used as a surrogate to study and understand the regulation, signaling, and metabolism of NO_2_-FA given its additional stability and well developed synthetic routes (Freeman et al., 2008). The isomers of NO_2_-OA used in our experiments have been extensively characterized and correspond to an equal proportion of the 9-NO_2_-OA and 10-NO_2_-OA species (Woodcock et al., 2013). Exogenous application of NO_2_-OA for functional studies has already been used and reported in plants. In Arabidopsis, exogenous application of NO_2_-OA regulates the expression of specific genes as demonstrated by qRT-PCR analysis (Mata Perez et al., 2016a), and, in tomato and Arabidopsis NO_2_-OA, functions as a signal that triggers ROS production via NADPH oxidase activation (Arruebarrena Di Palma et al., 2020). We acknowledge the limitations of studying NO_2_-OA in Arabidopsis root development, as it has not been detected in the free fatty acid fraction in Arabidopsis seedlings (Mata-Perez et al., 2016a). Nevertheless, the detection of free acid NO_2_-OA has been challenging as it rapidly adducts to thiol-containing proteins and glutathione, as demonstrated in mice plasma (Rudolph et al., 2010). This concept is further supported by studies in olives, where NO_2_-OA was found conjugated to proteins and not in the free fatty acid fraction (Fazzari et al., 2014). Recently, NO_2_-OA was detected in the free fatty acid fraction of Brassica seeds and seedlings, further supporting their formation and important role in plant physiology (Vollar et al., 2020). The discovery of the formation of NO_2_-Ln in Arabidopsis seeds and seedlings and the detection of NO2-OA in Brassica supports the concept of NO_2_-FA as a class of signaling species in plants (Mata Perez et al., 2016a; Vollar er al., 2020). It is in this context that our findings of the effects of NO_2_-OA on root development gain relevance, suggesting that NO_2_-FAs have common important formation pathways and roles in seeds and seedlings in different plant species.

Here we show that exogenous application of NO_2_-OA inhibits the growth of the primary root in Arabidopsis. The study of different root areas showed that cell size in the elongation-differentiation zone is not affected by NO_2_-OA. On the other hand, roots treated with NO_2_-OA showed a reduction in the size of the division zone. The total number of cortex cells and the total size of the meristematic area are reduced in roots treated with NO_2_-OA, indicating that the process of cell division is affected. The existing bibliography on the biological effects of Nitro Fatty acid in plants, suggest that they are acting as NO donors. In tomato cell suspensions, NO_2_-OA was unable to induce NO production (Arruebarrena Di Palma et al., 2020). Interestingly, under our experimental conditions, NO_2_-OA is not affecting endogenous NO levels in Arabidopsis seedlings, indication that the inhibition of primary root growth by NO_2_-OA does not involve a NO dependent signaling

Root growth is controlled by cell division and cell elongation rates. Both processes are strongly regulated by hormones, mainly auxins and cytokinins. Although auxins strongly control root growth, it has been shown that the interaction between these two signals is relevant for proper development (Dello loio et al., 2008). Auxins help to promote cell division and maintain the niche of stem cells and promote cytokinin biosynthesis, which consequently favors cell elongation and differentiation (Dello Ioio et al., 2008, Dello Ioio et al., 2007, Miyawaki K, et al., 2004). There is a strong relationship between auxin signaling and NO in root development. Mutants with low endogenous NO levels such as *nia1nia2* and *noa1* have lower auxin endogenous concentration (Sanz et al., 2014). However, NO inhibits the transport of auxins from the stem apical meristem to the root apical meristem (Fernandez Marcos et al., 2011). The inhibitory effect on root growth by NO is related to the inhibition of the signaling cascade caused by auxins, since the reduction in *DR5:GUS* reporter activity is seen in the root tip, as a consequence of lower levels of expression of PIN1 proteins (Fernandez-Marcos et al., 2011). Our results show that NO_2_-OA has an inhibitory effect on primary root growth and is not due to an unbalance in auxin or cytokinin levels. This further supports the idea that the effect of NO_2_-OA on root growth is different than that reported for NO donors. An RNAseq analysis of Arabidopsis cells treated with NO_2_-Ln, showed expression regulation of a large number of genes, but none of them related to signaling mediated by auxins or cytokinins (Mata Perez et al., 2016a). Consistently, our results show that NO_2_-OA does not affect auxins or cytokinins signaling.

Since the meristematic area is reduced, we studied the possibility that NO_2_-OA could affect the stem cell niche. To do so, we used the transcriptional *proPLT1:CFP, proPLT2:CFP* and the translational *proPLT1:PLT1-CFP* constructions considering PLT are transcription factors essential for QC identity and stem cell activity. The PLT genes are transcribed in response to auxin accumulation and are dependent on auxin response transcription factors. There is a correlation between PLT levels and root location, showing high levels in the division area, moderate in the transition zone and low levels in the elongation/differentiation side (Aida M, et al., 2004, Galinha et al., 2007). The expression levels of *PLT1* and *PLT2* are related to the size of the meristem. The double mutant *plt1-4*;*plt2-2* showed a reduction in the size of the meristem and short primary root. On the other hand, overexpression of *PLT2* showed an increase in the size of the meristem, particularly due to an increase in the number of cells (Aida et al 2004; Kornet et al., 2009; Mähönen et al., 2014). The expression of PLT is strongly related to cell division and the inhibition of differentiation since the expression of *PLT2* in epidermal cells inhibits the formation of root hairs (Mähönen et al., 2014). Although there is a strong correlation between *PLT 1* and *PLT2* and auxin levels (Galinha et al., 2007), there is no decrease of *DR5:GUS* reporter activity in the *plt1-4; plt2-2* double mutant (Aida et al., 2004; Kornet et al., 2009). Roots treated with NO_2_-OA had shorter meristem and a lower number of cells, but this was not due to altered levels in terms of expression or location of PLT1/2. In addition, we studied the localization of the QC transcription factor WOX5. WOX5 maintains stem cells in Arabidopsis roots (Sarkar et al., 2007). *WOX5* expression was no affected by NO_2_-OA treatment. Visualization of *PLT1*, *PLT2* and *WOX5* allowed us discard the effect of NO_2_-OA treatment on those transcription factors. Together these experiments suggest that the auxin and cytokinin balance and the transcription factors PLT1, PLT2 and WOX5, which are necessary for the development and maintenance of the stem cell niche, are not affected by NO_2_-OA treatments. Based on the observed results in which the division area is affected but not the elongation zone, QC or stem cells, we can conclude that NO_2_-OA has a selective effect on the process of cell division.

The *CyclinB1.1* gene codifies to a mitotic cyclin. The transcription of *CycB1.1* is associated with the G2 and M phase and is a marker for active cell division (Ferreira et al., 1994b). The construction of promoter region of *CycB1.1* fused with GUS is widely used to study particularly mitotic cell in different plant tissue (Colón-Carmona et al., 1999; Burssens et al., 2000). The results obtained with the *pCYCB1.1-GUS* confirm that NO_2_-OA inhibits cell division process in a dose-dependent manner.

Nitrolipids are weak electrophiles that can bind covalently to cystein or histidine residues of protein by Michael addition and modify stability or function of proteins (Batthyany et al., 2006; Baker et al., 2007). In animal few proteins have been described to be modified by NO_2_-FA, being most of them related to anti-inflammatory processes, such as the Nuclear factor (erythroid-derived 2) -like 2 (Nrf2), peroxisome proliferator-activated receptor-γ (PPARγ) and nuclear factor-kappa B (NF-kB) (Cui et al., 2006; Kansanen et al., 2011; Li et al., 2008). In plants, post-translational modifications mediated by NO_2_-FA were, so far, only reported for APX (Aranda-Caño et al., 2019). In the same review by Aranda-Caño, the authors mention that they identified a high number of nitroalkylated proteins that increase in cell cultures treated with NO_2_-Ln. It has been described that defects in the dynamics of the ACTIN cytoskeleton produce different phenotypes in root hairs and primary root (Gilliland et al., 2002, Yi et al., 2005, Kandasamy et al., 2009). Recent studies show that the *S*-Sulfhydration of the ACT2, ACT7 and ACT8 proteins modifies the dynamics of the cytoskeleton, decreasing the levels of the filamentous with respect to the globular form of the protein causing inhibition in the growth of the primary root as well as in the length of root hairs (Li et al., 2018). In the present paper we show that NO_2_-OA has physiological effects on root development and, specifically, in the process of cell division in the meristematic zone. Further research will be needed to elucidate whether NO_2_-OA interfere with cell division by affecting, directly, the order of actin or actin filaments, or through the regulation of other cell cycle regulating proteins.

## MATERIALS AND METHODS

### Chemicals and Reagents

OA was purchased from Nu-Chek Prep (Elysian, MN). NO_2_-OA and biotinylated NO_2_-OA were synthesized and purified as previously described (Woodcock el at., 2013; Bonacci et al., 2011; respectively).

### Plant material and growth conditions

Seeds from wild type Arabidopsis (*Arabidopsis thaliana* Col-0), *pCYCB1;1:GUS* (Colón-Carmona et al., 1999), *TCS:GFP* (Zucher et al., 2013), *DR5_pro_:GUS* (Ulmasov et al., 1997)*, PLT1_pro_:PLT1-YFP, PLT1_pro_:CFP,* and *PLT2_pro_:CFP* (Galinha et al., 2007), *CYCB1;1_pro_:GUS* (Colón-Carmona et al., 1999), *WOX5_pro_:GFP* (Sarkar et al., 2007) and *DII-Venus* (Brunoud et al., 2012) were surface sterilized in 35% sodium hypochlorite, stratified for 48 hours at 4°C in darkness. Seeds were germinated on vertically oriented plates containing 0.5X Murashige and Skoog (MS) salt mixture with Gamborg’s vitamins and 0.8% agar, and grown at 25°C using a 16-h-light/8-h-dark photoperiod.

### Seedling treatments

Five-days old seedlings were transferred onto plates containing Murashige and Skoog (MS) salt mixture with Gamborg’s vitamins and 0.8% agar, and grown at 25°C using a 16-h-light/8-h-dark photoperiod with OA, NO_2_-OA or non-treated for other five days.

### Measurement of primary root length, meristem length and cell size

Root length and meristem length were assessed in at least five independent experiments. Primary root length was measured using the software analysis package Fiji. For statistical, we used R software, we applied one-way ANOVA and Tukey’s multiple comparison test for the experiments.

To measure number of cortex cells, meristem length and cell size, seedlings were fixed in Hoyer’s solution for 30 min. The material was observed on a Zeiss Axioplan imaging 2 microscope under DIC optics. We used Poisson’s test for the experiment. Images were captured on an Axiocam HRC CCD camera (Zeiss) using the Axiovision program (version 4.2). Images analysis was performed using software package Fiji. The size of elongated cells were measured on two cells per root, on cells immediately before to the first root hair cell on 6 individuals per treatment.

### Nitric oxide detection on roots

Arabidopsis-treated roots were incubated in the presence of 10 μM of the fluorescent probe 4-aminomethyl-2’,7’-difluorofluorescein diacetate (DAF-FM DA) for 30 min. Roots were visualized under epifluorescence microscopy (Ex/Em wavelengths 495/515 nm).

### ß-Glucuronidase (GUS) expression

For GUS staining, Arabidopsis treated roots were incubated in GUS-staining buffer [5 mM EDTA, 0.1% Triton X-100, 5 mM K_4_Fe(CN)6, 0.5 mM K_3_Fe (CN)6], and 1 mg/mL X-Gluc (Rose Scientific) in 50 mM NaPi buffer, pH 7.0] for 3 h at 37°C. Then the tissue was cleared with Hoyer’s solution for 30 min. The material was observed on a Zeiss Axioplan imaging 2 microscope under DIC optics. Images were captured on an Axiocam HRC CCD camera (Zeiss) using the Axiovision program (version 4.2). GUS staining was measure using Fiji software (Béziat et al., 2017), we defined a ROI to analyze all images of the respective experiment as described by Feraru et al 2019.

### Confocal microscopy

Plant materials used in this study were previously described: PLT1pro:PLT1-YFP, PLT1pro:CFP, and PLT2pro:CFP (Galinha et al., 2007), CYCB1;1pro:GUS (Colón-Carmona et al., 1999), WOX5pro:GFP (Sarkar et al., 2007) and DII-Venus (Brunoud et al., 2012).

Arabidopsis roots from *pPLT1:CFP* y *pPLT2:CFP* transgenic plants were observed in confocal microscopy (Nikon Eclipse C1 Plus Ex/Em, 458/515).

Arabidopsis roots from *TCS:GFP* transgenic plants and WOX5pro:GFP were observed in confocal microscopy (Nikon Eclipse C1 Plus Ex/Em, 488/561) and quantified using Fiji software. We defined a ROI in the region that showed the most representative signal distribution. We used the same ROI (size and shape) to analyze all images of the respective experiment.

Arabidopsis root from PLT1pro:PLT1-YFP and DII-Venus were observed in confocal microscopy (Nikon Eclipse C1 Plus Ex/Em, 514/550). For DII-Venus marker we define two sections of the root, down QC is the calyptra zone, and up QC is the region above QC including QC, SCN (stem-cell niche) and a portion of meristem as indicated in Figure 4. We used the same ROI (size and shape) to analyze all images of the respective experiment.

PLT1pro:CFP, PLT1pro:PLT1-YFP and PLT2pro:CFP fluorescence was measure using Fiji software as described by Ercoli et al 2018.

## Data Analysis

We used R software for all data analysis. The most representative images are shown throughout the article

## Acknowledgments

We thanks to Ramiro Paris, Ramiro Rodriguez y Magdalena Vázquez for technical assistance on reporters analysis and for providing material for the DII-VENUS, WOX5, CYCB1:1 and PLT experiments.

## Notes

### Competing Interest Statement

The authors have declared no competing interest.

## REFERENCES

Aida M., Beis D., Heidstra R., Willemsen V., Blilou I., et al. (2004) The PLETHORA genes mediate patterning of the Arabidopsis root stem cell niche. Cell. 2004; 119:109–20.

Aranda-Caño L., Sánchez-Calvo B., Begara-Morales J. C., Chaki M., Mata-Pérez C.,Padilla, et al. (2019). Post-Translational Modification of Proteins Mediated by Nitro-Fatty Acids in Plants: Nitroalkylation. Plants, 8(4), 82. doi:10.3390/plants8040082

Arruebarrena Di Palma A, Di Fino LM, Salvatore SR, D’Ambrosio JM, García-Mata C, Schopfer FJ, Laxalt AM (2020). Nitro-oleic acid triggers ROS production via NADPH oxidase activation in plants: A pharmacological approach. J Plant Physiol.; 246-247:153128.

Baker PR, Schopfer FJ, Sweeney S, Freeman BA (2004) Red cell membrane and plasma linoleic acid nitration products: synthesis, clinical identification, and quantitation. Proc.Natl. Acad. Sci. U.S.A 101: 11577–11582

Baker LM, Baker PR, Golin-Bisello F, Schopfer FJ, Fink M, Woodcock SR, Branchaud BP, Radi R, Freeman BA (2007) Nitro-fatty acid reaction with glutathione and cysteine. Kinetic analysis of thiol alkylation by a Michael addition reaction. J. Biol. Chem. 282: 31085–31093

Batthyany C., Schopfer FJ., Baker PR., Duran R., Baker LM., Huang Y et al. (2006) Reversible post-translational modification of proteins by nitrated fatty acids in vivo. J. Biol. Chem. 281: 20450–20463

Benkova E., Hejatko J.. (2009). Hormone interactions at the root apical meristem. Plant Mol Biol. 69:383–96.

Béziat C., Kleine-Vehn J., Feraru E (2017) Histochemical staining of β-Glucuronidase and its spatial quantification. Methods Mol Biol, 73–80.

Bonacci G., Asciutto EK., Woodcock SR., Salvatore SR., Freeman BA., Schopfer FJ (2011) Gas-phase fragmentation analysis of nitrofatty acids. J Am Soc Mass Spectrom 22: 1534–1551

Brunoud, G., Wells, D., Oliva, M. et al. (2012) A novel sensor to map auxin response and distribution at high spatio-temporal resolution. Nature 482, 103–106

Burssens S., Himanen K., van de Cotte B., Beeckman T., Van Montagu M., Inzé D., et al. (2000) Expression of cell cycle regulatory genes and morphological alterations in response to salt stress in *Arabidopsis thaliana*. Planta 5: 632–40

Colón-Carmona A., You R., Haimovitch-Gal T., Doerner P (1999) Technical advance: spatio-temporal analysis of mitotic activity with a labile cyclin-GUS fusion protein. Plant J. 503–8.

Cui T., Schopfer FJ., Zhang J., Chen K., Ichikawa T., Baker PR., et al. (2006) Nitrated fatty acids: Endogenous anti-inflammatory signaling mediators. J Biol Chem 281: 35686–35698

Dello Ioio R., Linhares FS., Scacchi E., Casamitjana-Martinez E., Heidstra R., et al. (2007) Cytokinins determine Arabidopsis root-meristem size by controlling cell differentiation. Curr Biol. 17:678–82.

Dello Ioio R., Nakamura K., Moubayidin L., Perilli S., Taniguchi M., et al. (2008) A genetic framework for the control of cell division and differentiation in the root meristem. Science. 322:1380–84.

Dolan, L., Janmaat, K., Willemsen, V., Linstead, P., Poethig, S., Roberts, R., and Scheres, B. (1993). Cellular organisation of he Arabidopsis thaliana root. Development 119, 71–84

Ercoli F., Vena R., Goldy C., Palatnik JF., Rodriguez R (2018) Analysis of expression gradients of development regulators in *Arabidopsis thaliana* roots. Methods Mol Biol.

Fazzari M., Trostchansky A., Schopfer FJ., Salvatore SR., Sánchez-Calvo B., Vitturi D., et al. (2014). Olives and olive oil are sources of electrophilic fatty acid nitroalkenes. PloS one 9, e84884.

Ferreira PCG., Hemerly AS., de Almeida Engler J., Van Montagu M., Engler G., Inzé D (1994b) Developmental expression of the *Arabidopsis* cyclin gene *cyc1At*. Plant Cell 6: 1763–1774

Fernández-Marcos M., Sanz L., Lewis DR., Muday GK., Lorenzo O (2011) Nitric oxide causes root apical meristem defects and growth inhibition while reducing PIN-FORMED 1 (PIN1)-dependent acropetal auxin transport. PNAS 108, 18506–18511

Fernández-Marcos M., Sanz L., Lorenzo O (2012) Nitric oxide. An emerging regulator of cell elongation during primary root growth. Plant Signaling & Behavior 7:2, 196–200

Feraru E., FeraruaM I., Barbez E., Waidmann S., Sun L., Gaidora A., Kleine-Vehna J (2019) PILS6 is a temperature-sensitive regulator of nuclear auxin input and organ growth in Arabidopsis thaliana. PNAS 116, 3893–3898

Freeman BA., Baker PR., Schopfer FJ., Woodcock SR., Napolitano A., d’Ischia M (2008) Nitro-fatty acid formation and signaling. J Biol Chem 283: 15515–15519.

Galinha C., Hofhuis H., Luijten M., Willemsen V., Blilou I., et al. (2007) PLETHORA proteins as dose dependent master regulators of Arabidopsis root development. Nature. 449:1053–57.

Gilliland LU., Kandasamy MK., Pawloski LC., Meagher RB.. (2002) Both vegetative and reproductive actin isovariants complement the stunted root hair phenotype of the Arabidopsis act2-1 mutation. Plant Physiol. 130:2199–209

Grieneisen VA., Xu J., Marée AFM., Hogeweg P., Scheres B. (2007) Auxin transport is sufficient to generate a maximum and gradient guiding root growth. Nature. 449:1008–13.

Heyman, J., Kumpf, R.P., and De Veylder, L. (2014). A quiescent path to plantlongevity. Trends Cell Biol. 24, 443–448

Kandasamy MK., McKinney MC., Meagher RB (2009). A Single Vegetative Actin Isovariant Overexpressed under the Control of Multiple Regulatory Sequences Is Sufficient for Normal Arabidopsis Development Plant Cell. 21: 701–718

Kansanen E., Bonacci G., Schopfer FJ., Kuosmanen SM., Tong KI., Leinonen H., et al. (2011). Electrophilic nitro-fatty acids activate NRF2 by a KEAP1 cysteine 151-independent mechanism. Journal of Biological Chemistry 286, 14019–14027.

Kornet N., Scheres B. (2009) Members of the GCN5 histone acetyltransferase complex regulate PLETHORA-mediated root stem cell niche maintenance and transit amplifying cell proliferation in Arabidopsis. The Plant cell. 21:1070–1079.

Li Y., Zhang J., Schopfer FJ., Martynowski D., Garcia-Barrio MT., Kovach A., Suino-Powell K., et al. (2008) Molecular recognition of nitrated fatty acids by PPAR gamma. Nat Struct Mol Biol. 15:865–7.

Li J., Chen S., Wang X., Shi C., Liu H., Yang J., et al. (2018) Hydrogen Sulfide Disturbs Actin Polymerization via *S*-Sulfhydration Resulting in Stunted Root Hair Growth. Plant Physiol. 178:936–949.

Lombardo MC., Graziano M., Polacco JC., Lamattina L. (2006). Nitric oxide functions as a positive regulator of root hair development. Plant signaling & behavior 1, 28.

Mähönen AP., Ten Tusscher K., Siligato R., Smetana O., Díaz-Triviño S., Salojärvi J., et al. (2014). PLETHORA gradient formation mechanism separates auxin responses. Nature. 51:125–129.

Mata-Pérez C., Sánchez-Calvo B., Begara-Morales JC., Carreras A., Padilla MN., Melguizo M., et al. (2016c) Nitro-linolenic acid is a nitric oxide donor. Nitric Oxide - Biol Chem 57: 57–63

Mata-Pérez C., Sánchez-Calvo B., Begara-Morales JC., Padilla MN., Valderrama R., Corpas FJ., et al. (2016b) Nitric oxide release from nitro-fatty acids in Arabidopsis roots. Plant Signal Behav 11: 3–6

Mata-Pérez C., Sánchez-Calvo B., Padilla MN., Begara-Morales JC., Luque F., Melguizo M., et al. (2016a) Nitro-fatty acids in plant signaling: nitro-linolenic acid induces the molecular chaperone network in Arabidopsis. Plant Physiol 170: 686–701

Miyawaki K., Matsumoto-Kitano M., Kakimoto T. (2004). Expression of cytokinin biosynthetic isopentenyltransferase genes in Arabidopsis: tissue specificity and regulation by auxin, cytokinin, and nitrate. Plant J. 37:128–38.

Pagnussat GC., Lanteri ML., Lombardo MC., Lamattina L. (2004). Nitric oxide mediates the indole acetic acid induction activation of a mitogen-activated protein kinase cascade involved in adventitious root development. Plant Physiology 135, 279–286.

Perilli S., Sabatini S. (2010) Analysis of root meristem size development. Methods Mol Biol. 655:177–87.

Petricka JJ., Winter CM., Benfey PN. (2012) Control of Arabidopsis root development. Annu Rev Plant Biol. 2012;63:563–90

Rubbo H., Radi R., Trujillo M., Telleri R., Kalyanaraman B., Barnes S., et al. (1994) Nitric oxide regulation of superoxide and peroxynitrite-dependent lipid peroxidation. Formation of novel nitrogen-containing oxidized lipid derivatives. J Biol Chem 269: 26066–26075

Rubbo H., Radi R (2008) Protein and lipid nitration: role in redox signaling and injury. Biochim Biophys Acta 1780: 1318–1324

Rudolph TK., Rudolph V., Edreira MM., Cole MP, Bonacci G, Schopfer FJ, Woodcock SR, Franek A., Pekarova M., Khoo NKH, et al (2010) Nitro-fatty acids reduce atherosclerosis in apolipoprotein E-deficient mice. Arterioscler Thromb Vasc Biol 30: 938–945

Sabatini S., Heidstra R., Wildwater M., Scheres B. (2003) SCARECROW is involved in positioning the stem cell niche in the Arabidopsis root meristem. Genes Dev. 17:354–8.

Sanz L., Fernández-Marcos M., Modrego A., Lewis DR., Muday GK., Pollmann S., et al. (2014) Nitric oxide plays a role in stem cell niche homeostasis through its interaction with auxin. Plant Physiol. 166:1972–84

Sarkar, A., Luijten, M., Miyashima, S. et al. (2007). Conserved factors regulate signalling in *Arabidopsis thaliana* shoot and root stem cell organizers. Nature 446, 811–814

Scheres, B., Wolkenfelt, H., Willemsen, V., Terlouw, M., Lawson, E., Dean, C., and Weisbeek P (1994) Embryonic origin of the Arabidopsis primary root and root meristem initials. Development 120:2475–2487

Scheres B. (2007) Stem-cell niches: nursery rhymes across kingdoms. Nature Reviews. Molecular Cell Biology. 8:345–354.

Schopfer FJ., Cipollina C., Freeman BA (2011) Formation and signaling actions of electrophilic lipids, Chem. Rev. 111 5997–6021.

Trostchansky A., Rubbo H (2008) Nitrated fatty acids: mechanisms of formation, chemical characterization, and biological properties, Free Radic. Biol. Med. 44 1887–1896.

Ulmasov T., Murfett J., Hagen G., Guilfoyle TJ (1997) Aux/IAA proteins repress expression of reporter genes containing natural and highly active synthetic auxin response elements. Plant Cell. 1963–71.

van den Berg C., Willemsen V., Hendriks G., Weisbeek P., Scheres B. (1997) Short-range control of cell differentiation in the Arabidopsis root meristem. Nature. 390:287–89.

Vollár M., Feigl G., Oláh D., Horváth A,. Molnár. A, Kúsz N et al (2020) Nitro-Oleic Acid in Seeds and Di_erently Developed Seedlings of Brassica napus L. Plants 9

Weisbeek, P. (1994). Embryonic origin of the Arabidopsis primary root and root meristem. Development 120, 2475–2487

Wolters H., Jürgens G. (2009) Survival of the flexible: hormonal growth control and adaptation in plant development. Nat Rev Genet. 10:305–17

Woodcock SR., Bonacci G., Gelhaus SL., Schopfer FJ (2013) Nitrated fatty acids: synthesis and measurement. Free Radic Biol Med. 14–26

Xu J., Hofhuis H., Heidstra R., Sauer M., Friml J., Scheres B. (2006) A molecular framework for plant regeneration. Science. 311:385–8

Yi K., Guo C., Chen D., Zhao B., Yang B., Ren H. (2005) Cloning and functional characterization of a formin-like protein (AtFH8) from Arabidopsis. Plant Physiol. 138:1071–82.

Zürcher E., Tavor-Deslex D., Lituiev D., Enkerli K., Tarr P T., Müller B (2013) A Robust and Sensitive Synthetic Sensor to Monitor the Transcriptional Output of the Cytokinin Signaling Network in Planta Plant Physiology 161, 1066–1075

Zurcher E., Liu J., di Donato M., Geisler M., Muller B (2016): Plant development regulated by cytokinin sinks. Science 353:1027–1030.

