## Supplementary figures and images for "Exogenous Nitro-Oleic Acid inhibits primary root growth by reducing the mitosis in the meristem in *Arabidopsis thaliana*"

### Supplemental figure 1

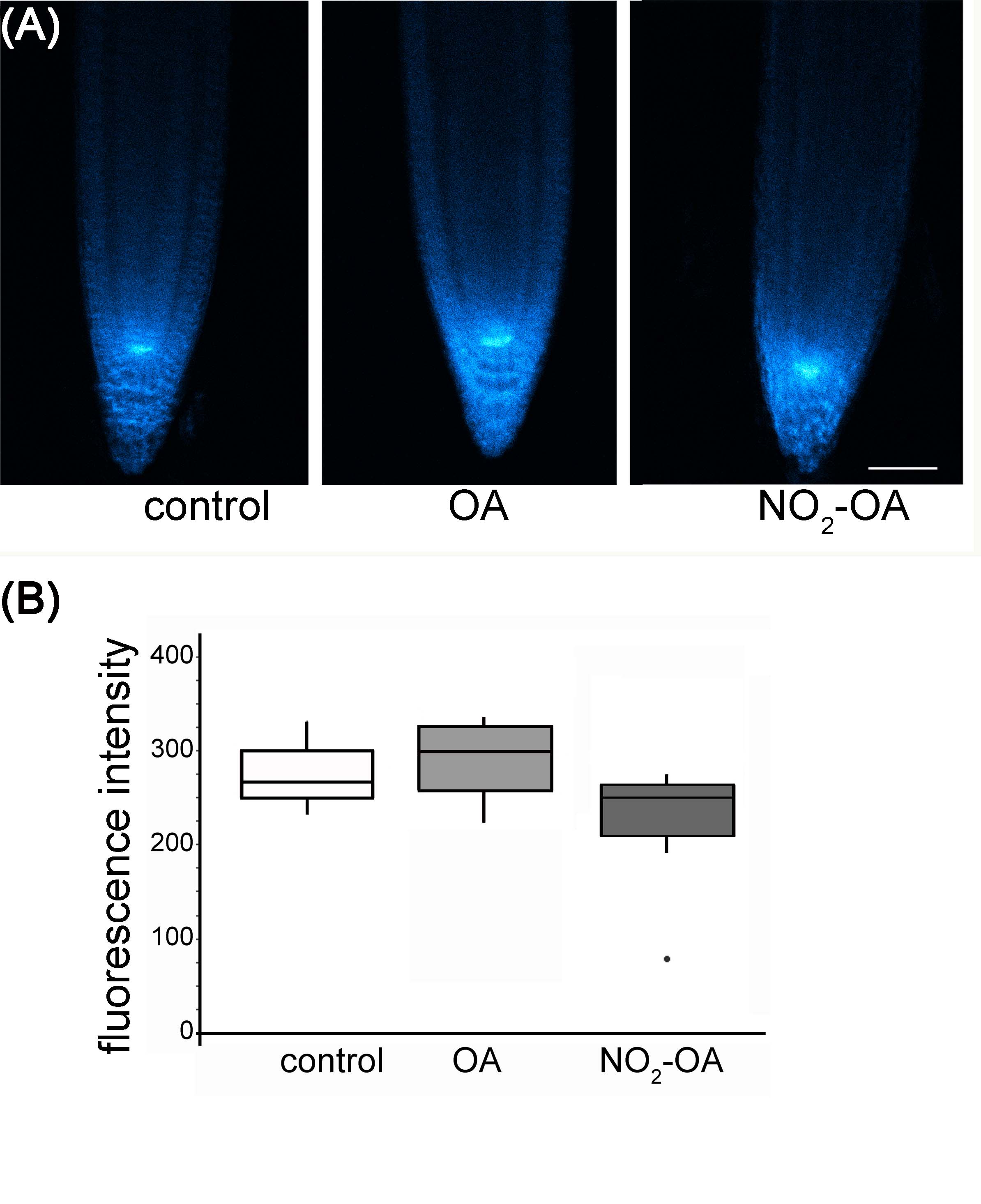
